# Cannabinoid signaling promotes the reprogramming of Muller glia into proliferating progenitor cells

**DOI:** 10.1101/2021.03.25.436969

**Authors:** Warren A. Campbell, Sydney Blum, Alana Reske, Thanh Hoang, Seth Blackshaw, Andy J. Fischer

**Affiliations:** Department of Neuroscience, College of Medicine, The Ohio State University, Columbus, OH; Solomon H. Snyder Department of Neuroscience, Johns Hopkins University School of Medicine, Baltimore, MD

**Keywords:** Endocannabinoids, Müller glia, Müller glia derived progenitor cells, scRNA-seq, Müller glia reprogramming

## Abstract

Endocannabinoids (eCB) are lipid-based neurotransmitters that are known to influence synaptic function in the visual system. eCBs are also known to suppress neuroinflammation in different pathological states. However, nothing is known about the roles of the eCB system during reprogramming of Müller glia (MG) into proliferating progenitor-like cells in the retina. Accordingly, we used the chick and mouse model to characterize expression patterns of eCB-related genes and applied pharmacological agents to examine how the eCB system impacts glial reactivity and the capacity of MG to become Müller glia-derived progenitor cells (MGPCs). We probed single cell RNA-seq libraries to identify eCB-related genes and identify cells with dynamic patterns of expression in damaged retinas. MG and inner retinal neurons expressed the eCB receptor *CNR1*, as well as enzymes involved in eCB metabolism. In the chick, intraocular injections of 2-Arachidonoylglycerol (2-AG) and Anandamide (AEA) potentiated the formation of MGPCs. Consistent with these findings, CNR1-agonists and MGLL-inhibitor promoted reprogramming, whereas CNR1-antagonist and inhibitors of eCB synthesis suppressed reprogramming. Surprisingly, retinal microglia were largely unaffected by increases or decreases in eCB signaling in both chick and mouse models. However, eCB-signaling suppressed the activation of NFkB-reporter in MG in damaged mouse retinas. We conclude that the eCB system in the retina influences the reactivity of MG and is important for regulating glial reactivity and the reprogramming of MG into proliferating MGPCs, but not for regulating the reactivity of immune cells in the retina.

**Main Points:** Müller glia express CNR1 receptor and endocannabinoid synthesis genes.

Endocannabinoids after retinal damage promote the formation of Müller glia derived progenitor cells in chick.

Endocannabinoids reduce NFkB activity in mouse Müller glia.

## Introduction

The endocannabinoid (eCB) system has been well-studied in the visual system and is known to modulate physiologic functions in different ocular tissues, including the retina (reviewed by (Schwitzer et al., 2016). The eCB system consists of cannabinoid receptors 1 and 2 (*CNR1*, *CNR2*), endogenous ligands 2-Arachidonoylglycerol (2-AG) and Arachidonoylethanolamide (AEA), and the enzymes that control ligand synthesis and degradation. The eCB pathway has been identified in the retinas of different vertebrates including embryonic chick (da Silva Sampaio et al., 2018), goldfish (Yazulla et al., 2000), rat (Yang et al., 2016), bovine (Bisogno et al., 1999), porcine (Matsuda et al., 1997), mouse (Hu et al., 2010), and human (Straiker et al., 1999). The expression of CNR1 and CNR2 receptors in the central nervous system varies across species but typically includes distinct types of neurons, astrocytes, microglia, and Müller glia. Activation of eCB receptors is known to modulate neurotransmission (Diana and Bregestovski, 2005), synaptic plasticity (Xu and Chen, 2015), neuroinflammation (Centonze et al., 2007), and neuroprotection (Slusar et al., 2013).

Müller glia (MG) are thought to play a role in regulating eCBs in the retina. Both CNR1 and CNR2 receptors have been identified in goldfish MG (Yazulla et al., 2000), and CNR2 receptors have been identified in the retinas of vervet monkeys (Bouskila et al., 2013). eCBs have been shown to modify activity or suppress T-type voltage gated calcium channels in rat MG (Yang et al., 2016) and modulate the inflammatory micro-environment (Silverman and Wong, 2018). MG possess pathogen- and damage-associated molecular pattern (PAMP/DAMP) receptors to respond to pathological conditions (Kumar and Shamsuddin, 2012; Kumar et al., 2013; Shamsuddin and Kumar, 2011). Activation leads to the secretion of pro-inflammatory cytokines to facilitate the migration and activation of macrophages and microglia (Inoue et al., 1996). At the same time, retinal microglia become reactive and coordinate inflammation with MG, which results in NF-kB activation, concomitant reactive gliosis, and formation of MGPCs (Palazzo et al., 2019). However, MG also produce anti-inflammatory signals such as TGFB2 (Palazzo et al., 2020) and TIMP3 (Campbell et al., 2019) to suppress inflammation. eCBs are believed to have anti-inflammatory actions within the central nervous system (Nagarkatti et al., 2009). Little is known about how eCBs influence inflammation in the retina and whether eCBs impact the ability of MG to reprogram into MG-derived progenitor cells (MGPCs).

The impact of inflammatory signals on MG is context specific, dependent on the combination of cytokines and the model of damage. In zebrafish, TNFa (Iribarne et al., 2019) and IL-6 (Zhao et al., 2014) are necessary for MG to transition to a reactive state into a proliferating progenitor-like cells. In the chick, by comparison, TNF alone does not induce MGPCs and activation of the NF-kB pathway inhibits the formation of MGPCs (Hoang et al., 2020; Palazzo et al., 2020). When microglia are ablated, MGPCs fail to form (Fischer et al., 2014), and the effects of NF-kB-inhibition are reversed to promote the formation of MGPCs (Palazzo et al., 2020). In damaged mouse retinas, reactive MG rapidly transition into a gliotic state and are forced back into a resting state, in part, by regulatory networks involving NF-kB-related factors (Hoang et al., 2020). This suggests that there is an important balance of inflammatory cytokines and timing of signals to drive the reprogramming of MG to dedifferentiate and proliferate as MGPCs. It is currently thought that rapid induction of microglial reactivity is required to “kick-start” MG reactivity as an initial step of reprogramming (Fischer et al., 2014; White et al., 2017), whereas sustained elevated microglial reactivity suppresses the neuronal differentiation of progeny produced by MGPCs (Palazzo et al., 2020; Todd et al., 2020).

In this study we investigate how eCBs influence glial reactivity, inflammation, and reprogramming of MG in the chick retina. Using scRNA-seq, we analyze the expression pattern of genes in the eCB system and changes in these genes following retinal damage. We apply pharmacological agents to activate or inhibit eCB-signaling and assess changes in glial activation and reprogramming of MG into proliferating MGPCs.

## Methods and Materials

### Animals

The animals approved in these experiments followed guidelines established by the National Institutes of Health and IACUC at The Ohio State University. P0 wildtype leghorn chicks (*Gallus gallus domesticus*) were obtained from Meyer Hatchery (Polk, Ohio). Post-hatch chicks were housed in stainless-steel brooders at 25°C with a diurnal cycle of 12 hours light, 12 hours dark (8:00 AM-8:00 PM) and provided water and Purina^tm^ chick starter *ad libitum*.

### Intraocular injections

Chicks were anesthetized with 2.5% isoflurane mixed with oxygen from a non-rebreathing vaporizer. The intraocular injections were performed as previously described (Fischer et al., 1998). With all injection paradigms, both pharmacological and vehicle treatments were administered to the right and left eye respectively. Compounds were injected in 20 μl sterile saline with 0.05 mg/ml bovine serum albumin added as a carrier. Compounds included: NMDA (500nmol dose high dose, 60nmol low dose; Sigma-Aldrich), JJKK048 (0.25mg/dose Sigma-Aldrich), ARN19874 (0.25mg/dose AOBIOUS), rimonabant (0.25mg/dose Sigma-Aldrich), PF 04457845 (0.25mg/dose Sigma-Aldrich), Orlistat (0.25mg/dose Sigma-Aldrich), URB 597 (0.25mg/dose Sigma-Aldrich). 5-Ethynyl-2’-deoxyuridine (EdU) was intravitreally injected to label the nuclei of proliferating cells. Injection paradigms are included in each figure.

### Enzyme-linked Immunosorbent Assay

Endocannabinoids were extracted from retinal tissue and screened for 2-AG levels using a direct competitive enzyme linked immunosorbent assay (MyBioSource). Three retinas were extracted from each treatment group and placed in 5:3 homogenization solution (formic acid pH = 3): extraction solution (9:1 ethylacetate:hexane) on ice. The tissue was homogenized with high intensity sonication on ice, frozen at −20, and the nonaqueous fraction was removed for evaporation and rehydration in DMSO. The lipid extract was applied to the wells of ELISA and the protocol was followed per the manufacturer’s instructions.

### Single Cell RNA sequencing of retinas

Retinas were obtained from postnatal chick and adult mice. Isolated retinas were dissociated in a 0.25% papain solution in Hank’s balanced salt solution (HBSS), pH = 7.4, for 30 minutes, and suspensions were frequently triturated. The dissociated cells were passed through a sterile 70µm filter to remove large particulate debris. Dissociated cells were assessed for viability (Countess II; Invitrogen) and cell-density diluted to 700 cell/µl. Each single cell cDNA library was prepared for a target of 10,000 cells per sample. The cell suspension and Chromium Single Cell 3’ V3 reagents (10X Genomics) were loaded onto chips to capture individual cells with individual gel beads in emulsion (GEMs) using 10X Chromium Controller. cDNA and library amplification for an optimal signal was 12 and 10 cycles respectively. Samples were multiplexed for sequencing on Illumina’s Novaseq6000 (Novogene). Sequencer files were converted from a BCL to a Fastq format, where the sequence files were de-multiplexed, aligned, and annotated using the chick ENSMBL database (GRCg6a, Ensembl release 94) and Cell Ranger software (10x Genomics). Using Seurat toolkits, Uniform Manifold Approximation and Projection for Dimension Reduction (UMAP) plots were generated from aggregates of multiple scRNA-seq libraries (Butler et al., 2018; Satija et al., 2015). Compiled in each UMAP plot are two biological library replicates for each experimental condition. Seurat was used to construct violin/scatter plots. Significance of difference in violin/scatter plots was determined using a Wilcoxon Rank Sum test with Bonferroni correction. Genes that were used to identify different types of retinal cells included the following: (1) Müller glia: *GLUL, VIM, SCL1A3, RLBP1*, (2) MGPCs: *PCNA, CDK1, TOP2A, ASCL1*, (3) microglia: *C1QA, C1QB, CCL4, CSF1R, TMEM22*, (4) ganglion cells: *THY1, POU4F2, RBPMS2, NEFL, NEFM*, (5) amacrine cells: *GAD67, CALB2, TFAP2A*, (6) horizontal cells: *PROX1, CALB2, NTRK1*, (7) bipolar cells: *VSX1, OTX2, GRIK1, GABRA1*, and (7) cone photoreceptors: *CALB1, GNAT2, OPN1LW*, and (8) rod photoreceptors: *RHO, NR2E3, ARR3.* scRNA-seq libraries can be queried at: https://proteinpaint.stjude.org/F/2019.retina.scRNA.html

### Fixation, sectioning, and immunocytochemistry

Ocular tissues were fixed, sectioned, and labeled via immunohistochemistry as described previously (Fischer et al., 2008, 2009a). Dilutions and commercial sources of antibodies used in this study are listed in table 2. Labeling was not due to non-specific labeling of secondary antibodies or tissue autofluorescence because sections incubated with secondary antibodies alone were devoid of fluorescence. Secondary antibodies included donkey-anti-goat-Alexa488/568, goat-anti-rabbit-Alexa488/568/647, goat-anti-mouse-Alexa488/568/647, goat-anti-rat-Alexa488 (Life Technologies) diluted to 1:1000 in PBS and 0.2% Triton X-100.

### Labeling for EdU

For the detection of nuclei that incorporated EdU, immunolabeled sections were fixed in 4% formaldehyde in 0.1M PBS pH 7.4 for 5 minutes at room temperature. Samples were washed for 5 minutes with PBS, permeabilized with 0.5% Triton X-100 in PBS for 1 minute at room temperature and washed twice for 5 minutes in PBS. Sections were incubated for 30 minutes at room temperature in a buffer consisting of 100 mM Tris, 8 mM CuSO_4_, and 100 mM ascorbic acid in dH_2_O. The Alexa Fluor 568 Azide (Thermo Fisher Scientific) was added to the buffer at a 1:100 dilution.

### Terminal deoxynucleotidyl transferase dUTP nick end labeling (TUNEL)

The TUNEL assay was implemented to identify dying cells by imaging fluorescent labeling of double stranded DNA breaks in nuclei. The *In Situ* Cell Death Kit (TMR red; Roche Applied Science) was applied to fixed retinal sections as per the manufacturer’s instructions.

### Photography, measurements, cell counts and statistics

Microscopy images of retinal sections were captured with the Leica DM5000B microscope with epifluorescence and the Leica DC500 digital camera. High resolution confocal images were obtained with a Leica SP8 available in The Department of Neuroscience Imaging Facility at The Ohio State University. Representative images are modified to have enhanced color, brightness, and contrast for improved clarity using Adobe Photoshop. In EdU proliferation assays, a fixed region of retina was counted and average numbers of Sox2 and EdU co-labeled cells. The retinal region selected for investigation was standardized between treatment and control groups to reduce variability and improve reproducibility.

Similar to previous reports (Fischer et al., 2009b, 2009c; Ghai et al., 2009), immunofluorescence was quantified by using Image J (NIH). Identical illumination, microscope, and camera settings were used to obtain images for quantification. Retinal areas were sampled from 5.4 MP digital images. These areas were randomly sampled over the inner nuclear layer (INL) where the nuclei of the bipolar and amacrine neurons were observed. Measurements of immunofluorescence were performed using ImagePro 6.2 as described previously (Ghai et al., 2009; Stanke et al., 2010; Todd and Fischer, 2015). The density sum was calculated as the total of pixel values for all pixels within thresholded regions. The mean density sum was calculated for the pixels within threshold regions from ≥5 retinas for each experimental condition. GraphPad Prism 6 was used for statistical analyses.

Measurements of immunofluorescence of CD45 in microglia were made from single optical confocal sections by selecting the total area of pixel values above threshold (≥70) for CD45 immunofluorescence. Measurements were made for regions containing pixels with intensity values of ≥70 or greater (0 = black and 255 = saturated). The total area was calculated for regions with pixel intensities above threshold. The intensity sum was calculated as the total of pixel values for all pixels within threshold regions. The mean intensity sum was calculated for the pixels within threshold regions from ≥5 retinas for each experimental condition. For characterization of the morphology of the individual microglia, a Sholl analysis was used to characterize the size, sphericity, and projections (ImageJ).

For statistical evaluation of differences across treatments, a two-tailed paired *t*-test was applied for intra-individual variability where each biological sample also served as its own control. For two treatment groups comparing inter-individual variability, a two-tailed unpaired *t*-test was applied. For multivariate analysis, an ANOVA with the associated Tukey Test was used to evaluate any significant differences between multiple groups.

## Results

### Patterns of expression of eCB-related genes

scRNA-seq libraries were aggregated from control retinas and retinas treated with NMDA-damage at different times (3, 12 and 48 hrs) after treatment. These libraries were clustered and analyzed for expression eCB-related genes under MG-reprogramming conditions (Fig. 1). UMAP plots were generated and the identity of clusters of cells established based on expression of cell-distinguishing markers (Fig. 1b,c). Resting MG occupied a discrete cluster of cells and expressed high levels of *GLUL, VIM* (Fig. 1d)*, RLBP1* and *CA2* (supplemental Fig. 1a-e,i). After damage, MG down-regulate these genes during transition to a reactive phenotype and up-regulate markers associated reactivity such as *MDK, HBEGF, MANF* (Fig. 1e, supplemental Fig. 1e,f,i), with some genes such as *TGFB2, ATF3* and *TNFRSF1A* upregulated with 3hrs of NMDA-treatment (supplemental Fig. 1f,g,i). Upregulation of progenitor- and proliferation-related genes was observed in MGPCs at 48hrs after NMDA-treatment (supplemental Fig. 1c,h,i), consistent with prior reports (Hoang et al., 2020; Campbell et al., 2021,).

**Figure 1.**
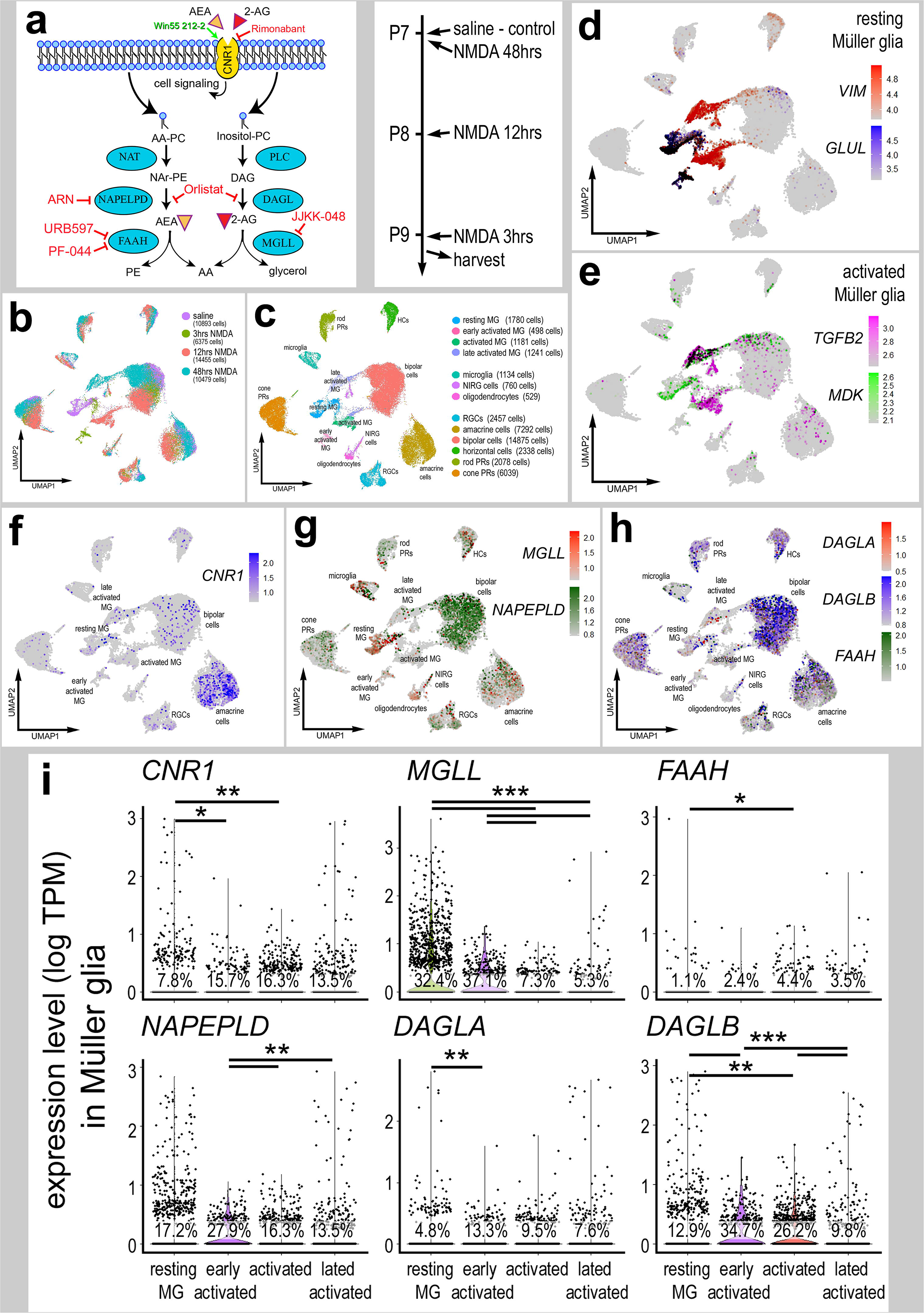
eCB-related genes are widely expressed in different types of retinal cells. Panel **a** illustrates a schematic diagram of the enzymes and receptors involved in eCB synthesis, degradation and signaling. scRNA-seq was used to identify patterns of expression of eCB-related genes among retinal cells. Patterns and levels of expression are presented in UMAP plots (**b**-**h**) and violin plots (**i**). scRNA-seq libraries were aggregated from control and treated 3hr, 12hr, and 48hr after NMDA-treatment (**b**). UMAP-ordered cells formed distinct clusters of neuronal cells, resting MG, early activated MG, activated MG and late activated MG (**c-e**). UMAP heatmaps of *CNR1*, *MGLL, NAPEPLD, DAGLA, DAGLB* and *FAAH* demonstrate patterns and levels of expression across different retinal cells, with black dots representing cells with expression of 2 or more genes (**f**-**h**). Violin plots illustrate relative levels and percent of expression in resting and activated MG (**i**). Violin plots illustrate levels of gene expression and significant changes (*p<0.01, **p<10exp-10, ***p<10exp-20) in levels that were determined by using a Wilcox rank sum with Bonferroni correction.

The expression of eCB-related genes in MG has been previously reported in developing chick retina (da Silva Sampaio et al., 2018). The eCB system includes receptors *CNR1* and *CNR2* and enzymes involved in the synthesis (*NAPEPLD, DAGLA* and *DAGLB*) and degradation (*FAAH* and *MGLL*) of 2-AG and AEA (Fig. 1a). We detected *CNR1*, *MGLL*, *DAGLA, DAGLB, NAPELPD and FAAH* in control retinas and at different times after NMDA-treatment (Fig. 1f-i). *CNR2* was not detected. *MGLL* was prevalent and highly expressed by resting MG, but down-regulated in activated MG (Fig. 1g,f). By comparison, levels of expression and prevalence of *CNR1* was high in many amacrine cells, and in a few ganglion and bipolar cells (Fig. 1f). *MGLL* and *NAPEPLD* were detected at high levels in many microglia, NIRG cells and bipolar cells, and in relatively few photoreceptors, ganglion and horizontal cells (Fig. 1e). *DAGLA* and *FAAH* had scattered expression across many retinal cell types, whereas *DAGLB* was prominently expressed at high levels in photoreceptors and inner retinal neurons (Fig. 1h). *CNR1* and eCB-related genes, except *FAAH*, were uniformly down-regulated in levels, but increased in prevalence in activated MG after NMDA-treatment (Fig. 1g-i).

To directly compare expression levels in MG across different treatment paradigms we isolated and re-aggregated MG from different treatment groups, including retinas treated with the combination of NMDA+insulin+FGF2, NMDA alone and insulin+FGF2 alone. Resting MG, activated MG from 24 hrs after NMDA-treatment and 2 doses of insulin+FGF2 formed distinct clusters of cells (Fig. 2a,b). Further, MGPCs formed discrete regions of cells wherein cell cycle progression formed the basis of spatial segregation with the majority of cells progressing through the cell cycle from retinas at 72 hrs after NMDA or NMDA+FGF2+insulin treatment (Fig. 2c,e). The largest increase in levels and prevalence of expression of *CNR1* was observed in reactive MG from 48+72hrs after NMDA (Fig. 2f,g). By comparison, the largest decrease in levels and prevalence of expression of *MGLL, DAGLA, DAGLB* and *NAPEPLD* were observed for MG clustered among treatments with insulin+FGF2 and in the MGPC3 cluster (Fig. 2f,g). Collectively, these findings suggest that the expression of *CNR1* by MG is changed in response to neuronal damage, whereas levels of *MGLL* and other eCB-related genes were down-regulated by treatment with NMDA (neuronal damage) or insulin and FGF2 (no neuronal damage).

**Figure 2.**
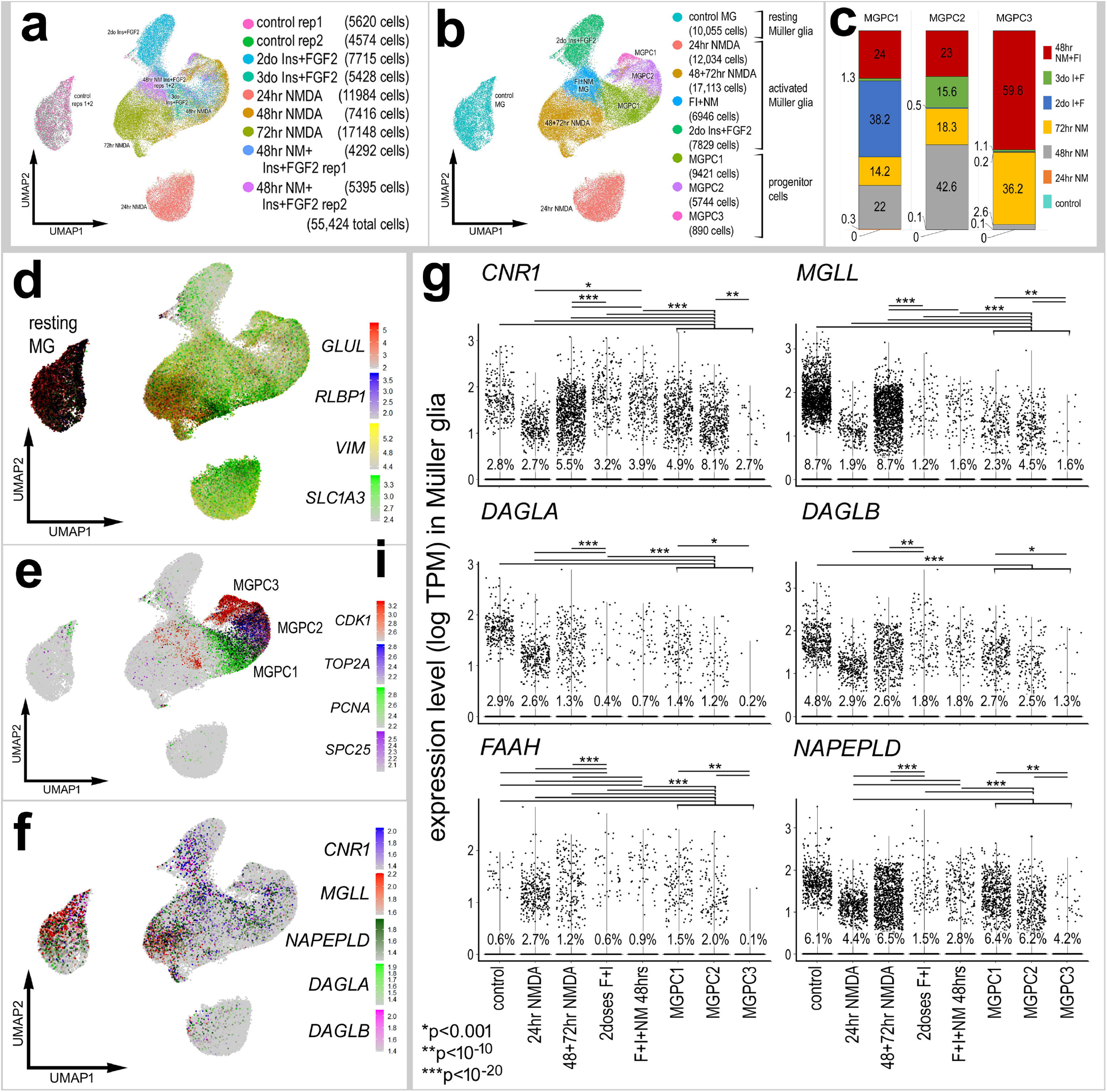
eCB-related genes are dynamically expressed by MG in response to damage or growth-factor treatment. scRNA-seq was used to identify patterns of expression of eCB genes in MG at several time points after NMDA damage or FGF + insulin growth factor treatment to form MGPCs. UMAP-clusters of MG were identified by expression of hallmark genes (**a,b,d**). Progenitors were then classified by different cell cycle and progenitor markers (**c, e, f**). Each dot represents one cell and black dots indicate cells with 2 or more genes expressed. The expression of eCB related genes was illustrated in a colored heatmap and in a violin plot violin plot with population percentages and statistical comparisons. (**g,h,i**). Significant difference (*p<0.01, **p<0.0001, ***p<<0.0001) was determined by using a Wilcox rank sum with Bonferroni correction. MG – Müller glia, MGPC – Müller glia-derived progenitor cell.

### eCBs promote the formation of MGPCs after damage

Although patterns of gene expression can be complex and context dependent, dynamic changes in mRNA levels are strongly correlated with changes in protein levels and function (Liu et al., 2016). Accordingly, we tested activation of eCB-signaling influence glial reactivity, neuronal survival and the formation of MGPCs. The ligand binding affinity of chick CNR receptors remains uncertain. Thus, 2-AG and AEA were co-injected to maximize the probability of activation of CNR1 receptors. We tested whether co-injection of 2-AG and AEA influenced the formation of proliferating MGPCs. Compared to numbers of proliferating MGPCs in NMDA-damaged retinas, treatment with eCBs resulted in a significant increase in numbers of Sox2/EdU-positive MGPCs (Fig. 3a,b). Consistent with these findings, numbers of proliferating MGPCs that expressed neurofilament and phospho-histone H3 (pHH3) were significantly increased by treatment with 2-AG and AEA (Fig. 3c,d). Levels of retinal damage influence the reprogramming of MG; there is a positive correlation between numbers of dying cells and numbers of proliferating MGPCs (Fischer and Reh, 2001; Fischer et al., 2004). Accordingly, we probed for numbers of dying cells by labeling for fragmented DNA using the TUNEL method. The number of TUNEL-positive cells was unchanged by 2-AG and AEA, suggesting that levels of cell death in NMDA-damaged retinas were unaffected by addition of eCBs (Fig 3e,f).

**Figure 3.**
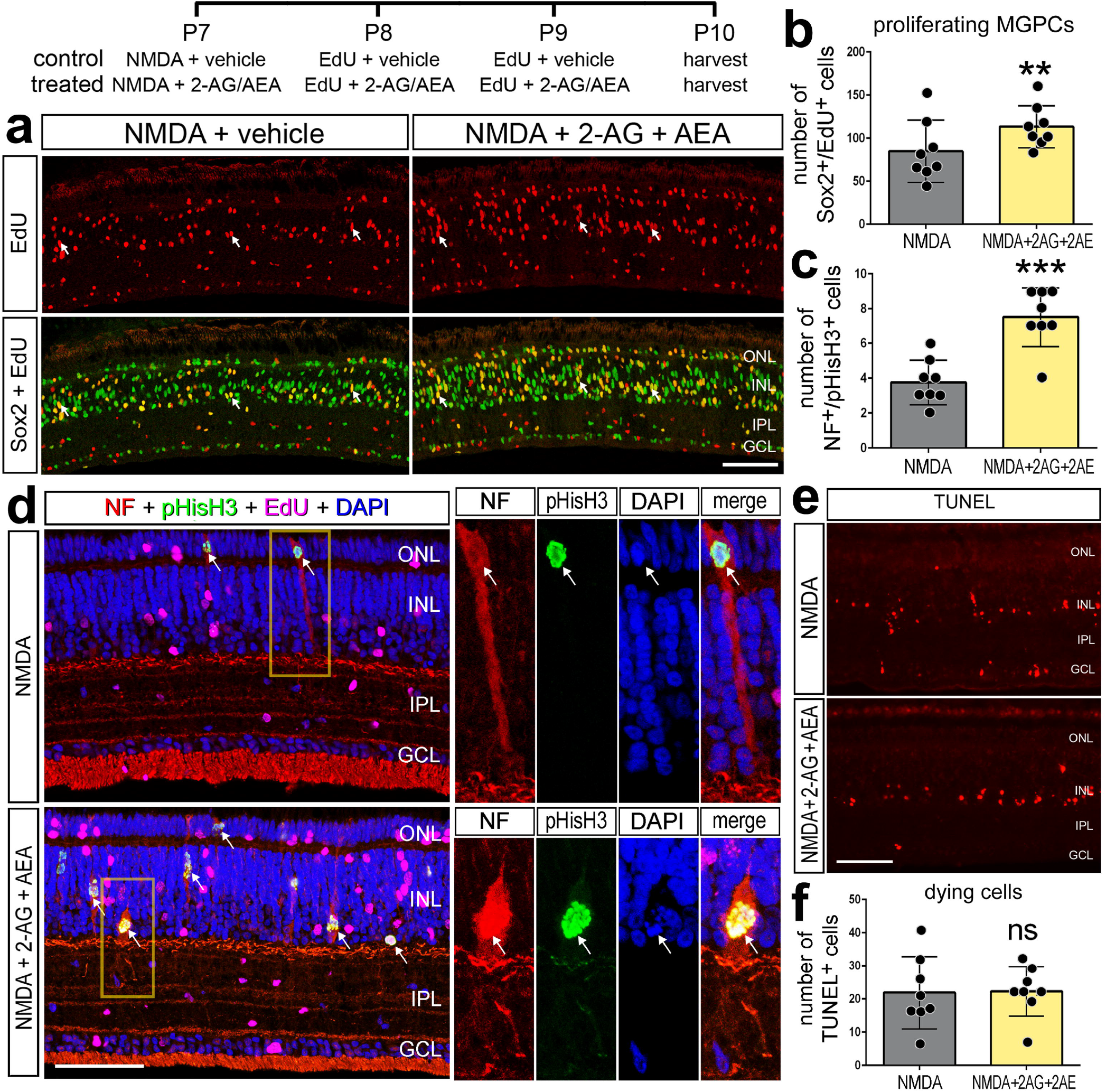
eCB increase numbers of proliferating MGPCs in damaged retinas. Chick eyes were injected with NMDA, AEA, 2-AG, and EdU according to the paradigm at the top figure. Eyes were harvested at 24 hrs after the last injection and retinas processed for immunolabeling. Retinas were labeled for Sox2 (green) and EdU^+^ (red) cells (**a**), neurofilament (red), phospho-Histone H3 (pHisH3, green), and DAPI (blue; **d**). Dying cells were labeled using the TUNEL assay (**e**). Histograms illustrate the mean (± SD) and each dot represents one biological replicate. Significance of difference (**p<0.01, ***p<0.001) was determined by using a paired *t*-test. Arrows indicate the nuclei of MG. The calibration bar panels **a, d** and **e** represent 50 µm. Abbreviations: ONL – outer nuclear layer, INL – inner nuclear layer, IPL – inner plexiform layer, GCL – ganglion cell layer, NF – neurofilament, ns – not significant.

### Targeting the eCB synthesis and degradation influences MG reprogramming

Since expression levels of eCB-related genes were changed in NMDA-damaged retinas, we investigated whether levels of eCBs were influenced by damage or drugs that interfere with synthesis or degradation of AEA and 2-AG. We applied Orlistat, an inhibitor of DAGL, to reduce eCB synthesis and JJKK-048, an inhibitor to MGLL, to suppress eCB degradation (Hillard, 2015). By using competitive inhibition ELISAs, we measured levels 2-AG and AEA in retinas treated with NMDA and inhibitors. We detected low levels of 2-AG in the retina, that did not significantly change with NMDA damage at 72 hours (Fig. 4a). Although we failed to detect a significant change in 2-AG with Orlistat treatment, injections of JJKK-048 resulted in a significant increase in retinal levels of 2-AG (Fig. 4a). AEA was not detectable within the threshold range of the ELISA; thus, inhibitor treatments had no detectable impact on levels of AEA (Fig. 4b). Since levels of AEA fell below levels of detection we did not probe for changes in AEA-levels following treatment with inhibitors of NAPEPLD or FAAH.

**Figure 4.**
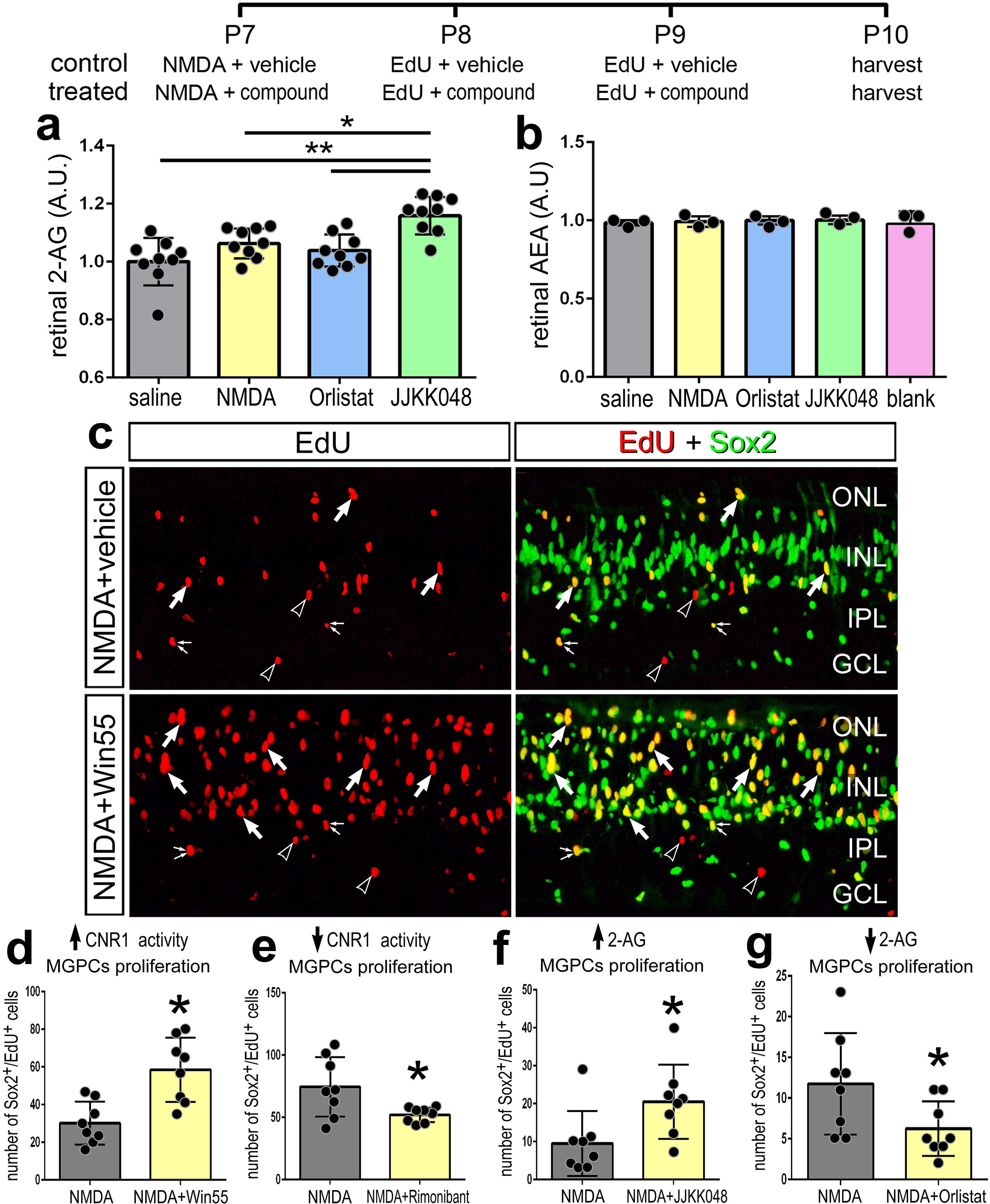
Small molecule drug targeting CNR1, DAGL and MGLL influence the formation of MGPCs. The treatment paradigm is illustrated at the top of the figure. Compounds included Orlistat (DAGL inhibitor), JJKK048 (MGLL inhibitor), Win55 (CNR1 agonist), and Rimonabant (CNR1 antagonist). Competitive inhibitor ELISAs of illustrate relative levels of 2-AG and AEA after NMDA damage and treatment with Orlistat or JJKK048 (**a,b**). Retinas were labeled for Sox2 (green) and EdU (red; **a**). Arrows indicate EdU+/Sox2+ nuclei of MGPCs, small double-arrows indicate EdU+/Sox2+ nuclei of NIRG cells in the IPL, and hollow arrow-heads indicate EdU+/Sox- nuclei of presumptive microglia. The histograms in **a,b** and **d-g** represents the mean (± SD) and each dot represents one biological replicate retina. The calibration bar in **c** represents 50 µm. Abbreviations: ONL – outer nuclear layer, INL – inner nuclear layer, IPL – inner plexiform layer, GCL – ganglion cell layer.

We next tested whether inhibition of enzymes that produce or degrade eCBs influence glial reactivity, cell death and the formation of MGPCs. We also targeted the CNR1 receptor with a small molecule agonist and an antagonist. Win-55, 212-2 (Win55) is a potent CNR agonist in humans, mice and chickens (Stincic and Hyson, 2011). Rimonabant is a potent and selective antagonist that inhibits CNR1-mediated cell-signaling (Ádám et al., 2008) (Hillard, 2015). Activation of CNR1 with Win55 increased numbers of proliferating MGPCs, whereas inhibition of CNR1 with rimonabant had the opposite effect (Fig. 4c,d,e). MGLL inhibitor (JJKK048), which increased levels of 2-AG (Fig. 4a), increased numbers of proliferating of MGPCs (Fig. 4f). By comparison, the DAGL inhibitor Orlistat significantly decreased numbers of MGPCs (Fig 4g). Overall, treatments expected to increase eCB-signaling increased MG reprogramming and treatments to decrease eCB-signaling decreased MG reprogramming.

We next targeted enzymes that influence the synthesis (NAPEPLD) or degradation (FAAH) of AEA. Inhibition of NAPELPD with ARN19784 had no effect upon numbers of proliferating MGPCs (Fig. 5a,b), whereas numbers of proliferating microglia were increased (Fig 5c,d) and numbers of proliferating NIRG cells and dying cells were decreased (Fig. 5e-h). By comparison, inhibition of FAAH with URB597 or PF-044 had no significant effect upon proliferating MGPCs, microglia and NIRG cells, or cell death (Fig. 5d,f,h). We bioinformatically isolated scRNA-seq data for microglia and performed a fine-grain analysis. Microglia formed discrete UMAP clustering of resting and activated microglia from control retinas and retinas at 3, 12 and 48 hrs after NMDA-treatment (Fig. 5i-l). We detected scattered expression of relatively high levels of *MGLL* and *NAPEPLD*, but no expression of *CNR1* (Fig. 5m) or *FAAH* (not shown).

**Figure 5.**
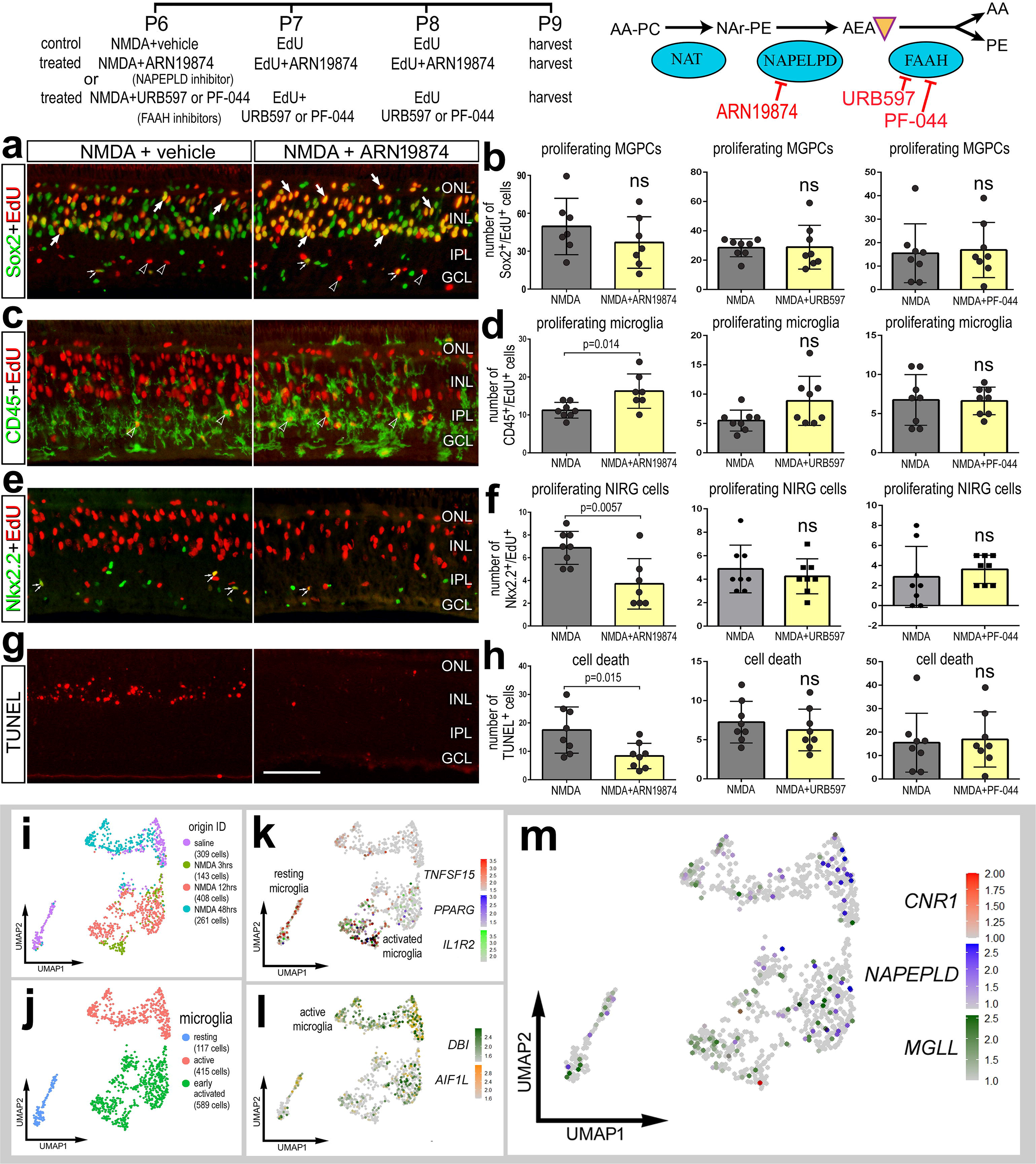
Targeting the AEA pathway does not influence the formation of MGPCs. The treatment paradigm is illustrated at the top of the figure. Compounds included ARN19874 (NAPEPLD inhibitor), URB597 (FAAH inhibitor) and PF-044 (FAAH inhibitor). Eyes were harvested at 24 hrs after the last injection and retinas processed for immunofluorescence. Retinal sections were labeled for Sox2 (green) and EdU (red; **a**), CD45 (green) and EdU (red; **c**), Nkx2.2 (green) and EdU (red, **e**), and cell death (TUNEL, red; **g**). The histograms in **b,d**,**f,h** represents the mean (± SD) number of proliferating MGPCs (**b**), proliferating microglia (**d**), proliferating NIRG cells (**f**), and dying cells (**h**). Each dot represents one biological replicate retina. Arrows indicate EdU+/Sox2+ nuclei of MGPCs, small double-arrows indicate EdU+/Nkx2.2+ nuclei of NIRG cells in the IPL, and hollow arrow-heads indicate EdU+/CD45+ nuclei of microglia. The calibration bar in **g** represents 50 µm and applies to panels **a,c,e** and **g**. Abbreviations: ONL – outer nuclear layer, INL – inner nuclear layer, IPL – inner plexiform layer, GCL – ganglion cell layer. Microglia were from aggregate scRNA-seq libraries were re-embedded and ordered in UMAP (**i**). Microglia from saline- and NMDA- treated retinas were clustered into resting, active and early activated cells (**j**). Clusters of cells were arranged according to markers such as *TNFSF15, PPARG, IL1R2, DBI* and *AIR1L* (**k,l**). *MGLL* and *NAPEPLD* had scattered expression across microglia in different clusters, whereas *CNR1* was not widely expressed (**m**).

Collectively, these data suggests that cells in chick retina support production of 2-AG over AEA in the context of damage and reprogramming. Further the reactivity of some microglia and NIRG cells, but not MG, is influenced by inhibition of NAPEPLD, and these responses are consistent with patterns of expression seen in scRNA-seq databases.

### Microglia Reactivity and eCBs

Retinal microglia serve homeostatic functions and mediate inflammation in response to damage and pathogens (Silverman and Wong, 2018). In response to excitotoxic damage in the chick, the microglia become reactive, leading to accumulation of monocytes, proliferation, and upregulation of inflammatory cytokines (Fischer et al., 2014). Given the known association of microglia, inflammation and eCB-signaling (Stella, 2009) and the dependence of MGPC formation on signals provided by reactive microglial (Fischer et al., 2014; Palazzo et al., 2020), we investigated the impact of drugs targeting CNR1 and 2-AG metabolism on microglia reactivity, proliferation and reactivity.

Microglia are sparsely distributed and highly ramified when quiescent, but become reactive and transiently accumulate after NMDA-treatment (Fischer et al., 2014). We applied established metrics of microglia reactivity in the chick model (Gallina, 2015), including microglia infiltration/accumulation, proliferation, CD45-intensity, cell area, and ramification were compared in different eCB targeted treatments. eCBs and small molecule inhibitors had no significant effect on the reactivity of microglia in damaged retinas (Fig. 6). Both the small molecule drugs and 2-AG/AEA did not influence total numbers of CD45^+^ cells compared to damage alone (Fig. 6a-c). Similarly, the number of proliferating CD45^+^ cells was unaffected by eCB treatments in damaged retinas (Fig. 5a-c). Similarly, the area and intensity of CD45^+^ immunolabeling were unaffected by drugs targeting eCBs (Fig. 6c). Using a Sholl analysis to quantify microglia shape, we quantified the maximum intersections (ramification index), mean intersections (centroid value), and maximum intersection radius (processes distribution). Although the morphology of resting microglia in saline treated retinas was significantly different from the morphology of microglia in NMDA-damaged retinas (Fig. 6d), the morphology of reactive microglia was unchanged by drugs targeting eCB receptors or metabolic enzymes (table 1).

**Figure 6.**
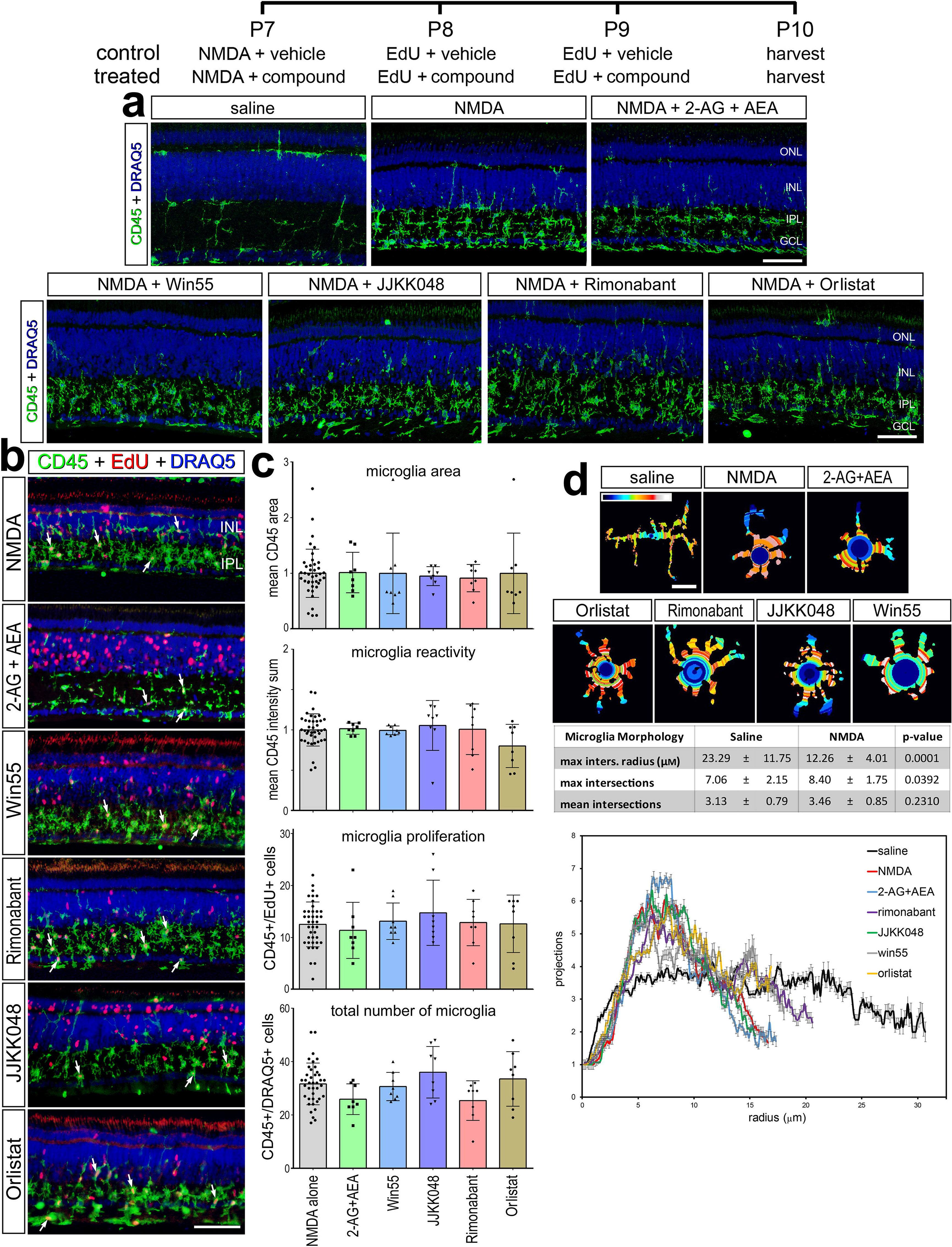
Microglia reactivity in damaged retina treated with eCBs. The treatment paradigm is illustrated at the top of the figure. Compounds included 2-AG+AEA (CNR1 agonists), Orlistat (DAGL inhibitor), JJKK048 (MGLL inhibitor), Win55 (CNR1 agonist), and Rimonabant (CNR1 antagonist). Eyes were harvested 24hrs after the last injection and retinas were processed for immunofluorescence. Retinal sections were labeled for CD45 (green) and DAPI (blue; **a**), or CD45 (green), EdU (red) and DAPI (blue; **b**). Microglial reactivity was assessed by measuring CD45 area and intensity, proliferation (EdU+), and total number of microglia (CD45+/DRAQ5+) (**b**). Arrows indicate the nuclei of microglia. Histograms in **c** illustrate the mean (±SD, control n = 40, treatment n = 8) and each symbol represents one biological replicate. The shape of the microglia was assessed using a Sholl analysis. Representative microglia from each condition is shown, with a heat map of radial intersections (blue = low, red/white = high) (**c**). The graph in **d** illustrates of the number of processes radially (µm±SE) from microglial nuclei from control (n=15) and NMDA damaged (n = 25) retinas. (**d**) Significance of difference was determined by using a one way ANOVA with corresponding Tukey’s test. The calibration bars panels **a, b** represent 50 µm, and the bar in d represents 5 µM. Abbreviations: ONL – outer nuclear layer, INL – inner nuclear layer, IPL – inner plexiform layer, GCL – ganglion cell layer.

**Table 1.** Sholl Analysis of microglia from the chick retina after damage and eCB treatment. Statistical analysis was performed with a one-way ANOVA and Tukey’s test (NMDA n = 25, treatments n = 15)

|  | vehicle | enhance signal |  |  | dampen signal |  | ANOVA |
| --- | --- | --- | --- | --- | --- | --- | --- |
| Microglia Morphology | NMDA | 2AG + AEA | JJKK-048 | Win55-212,2 | Rimonabant | Orlistat | p-value |
| max inters. radius (μM) | 12.26 ± 4.01 | 11.90 ± 3.72 | 11.99 ± 3.51 | 12.65 ± 5.44 | 14.01 ± 5.83 | 12.27 ± 2.96 | 0.782 |
| mean intersections | 8.40 ± 1.75 | 7.93 ± 1.83 | 8.80 ± 2.54 | 7.40 ± 1.76 | 8.20 ± 1.85 | 9.13 ± 1.88 | 0.189 |
| max intersections | 3.46 ± 0.85 | 3.36 ± 0.61 | 3.58 ± 0.86 | 3.44 ± 0.81 | 3.63 ± 0.70 | 0.36 ± 0.68 | 0.921 |

### NF-kB activation is reduced in mouse MG when promoting eCB signaling

NF-kB is a transcription factor known to be a primary transductor of the innate and adaptive immunity, and a central mediator of inflammatory responses to pathogens or tissue damage (Liu et al., 2017). In the chick, activation or inhibition of the NF-kB pathway has a significant impact on the ability of MG to become proliferating MGPCs (Palazzo et al., 2020). In the mouse retina, NF-kB-signaling has been implicated a signaling “hub” that may act to drive MG into a reactive state and then back into a resting state (Hoang et al., 2020). Accordingly, we investigated the anti-inflammatory properties of eCBs using the NF-kB reporter in the mouse retina. We used the mouse model because there are no cell-level read-outs of NF-kB-signaling available in the chick (Palazzo et al., 2020).

We first assessed the patterns of expression of eCB-related factors in normal and NMDA-damaged mouse retinas in aggregated scRNA-seq libraries. UMAP analysis of cells from control and damaged retinas revealed discrete clusters of cell types (Fig. 7a). Neurons from control and damaged retinas were clustered together regardless of time after NMDA-treatment (Fig. 7a). By contrast, resting MG, including MG from 48 to 72 hr after NMDA, and activated MG from 3, 6, 12, and 24 hr after treatment were spatially separated across the UMAP plot (Fig. 7c). Consistent with previous reports (Bouchard et al., 2015), *Cnr1* was detected in amacrine and ganglion cells (Fig. 7d), whereas *Cnr2* was not detected at significant levels in any retinal cells (not shown). By comparison, *Daglb* was detected in many retinal neurons including photoreceptors and bipolar cells, and *Mgll* was detected prominently in ganglion cells, glycinergic amacrine cells, and resting MG (Fig. 7e), similar to patterns seen in chick retinas (Fig. 1). *Dagla* was not detected (not shown). Although *Napepld* was not widely expressed, *Faah* had scattered expression in bipolar cells, and rod and cone photoreceptors (Fig. 7f).

**Figure 7.**
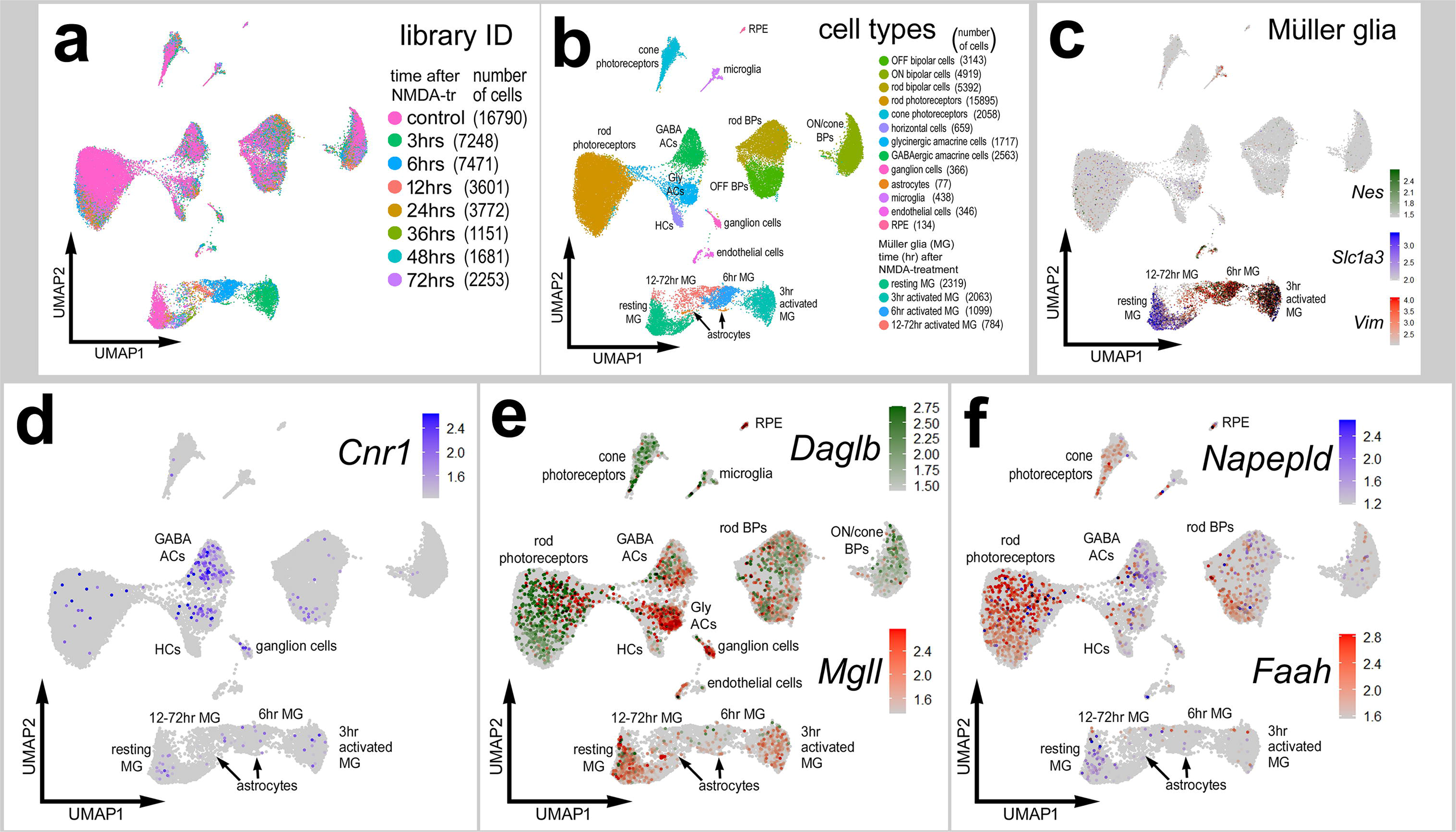
Patterns of expression of eCB-related genes in normal and NMDA- damaged mouse retinas. Cells were obtained from control retinas and from retinas at 3, 6, 12, 24, 36, 48 and 72hrs after NMDA-treatment and clustered in UMAP plots with each dot representing an individual cell (**a**). UMAP plots revealed distinct clustering of different types of retinal cells; resting MG (a mix of control, 48hr and 72hr NMDA-tr), 12-72 hr NMDA-tr MG (activated MG in violin plots), 6hrs NMDA-tr MG, 3hrs NMDA-tr MG, microglia, astrocytes, RPE cells, endothelial cells, retinal ganglion cells, horizontal cells (HCs), amacrine cells (ACs), bipolar cells (BPs), rod photoreceptors, and cone photoreceptors (**b**). Resting and activated MG were identified based on patterns of expression of *Slc1a3, Nes* and *Vim* (**c**). Cells were colored with a heatmap of expression of Cnr1*, Daglb, Mgll, Napepld and Faah* expression (**d**-**f**). Black dots indicate cells that express two or more markers.

We utilized the cis-NF-kB^eGFP^ reporter mouse line to visualize cells where p65 is driving transcription as a read-out of NF-kB-signaling (Magness et al., 2004). In undamaged retinas, NFkB reporter was observed in a few endothelial cell whereas eGFP reporter was not detected in any retinal neurons or glia (Fig. 8a). At 48hrs after NMDA damage significant numbers of MG express NFkB-eGFP (Fig. 8a,b). Treatment with CNR1 agonist Win55 or eCBs (2-AG/AEA) resulted in a significant reduction in numbers of MG that were eGFP-positive (Fig. 8a,c). By contrast, treatment with CNR1 antagonist (Rimonabant) significantly increased numbes of eGFP-positive MG (Fig. 8a,c). To determine if changes in cell death were influenced eCBs we performed TUNEL staining. There was no change in cell death in retinas treated with eCBs, Rimonabant or Win55 (Fig. 8d,e). In addition, there was there was no obvious change in microglial morphology (Fig. 8f) and no significant change in the accumulation of microglia in damaged retinas treated with eCBs, Rimonabant or Win55 (Fig. 8g). These data suggest that eCBs influence NF-kB signaling in mammalian MG, whereas neuroprotection and microglial reactivity are unaffected.

**Figure 8.**
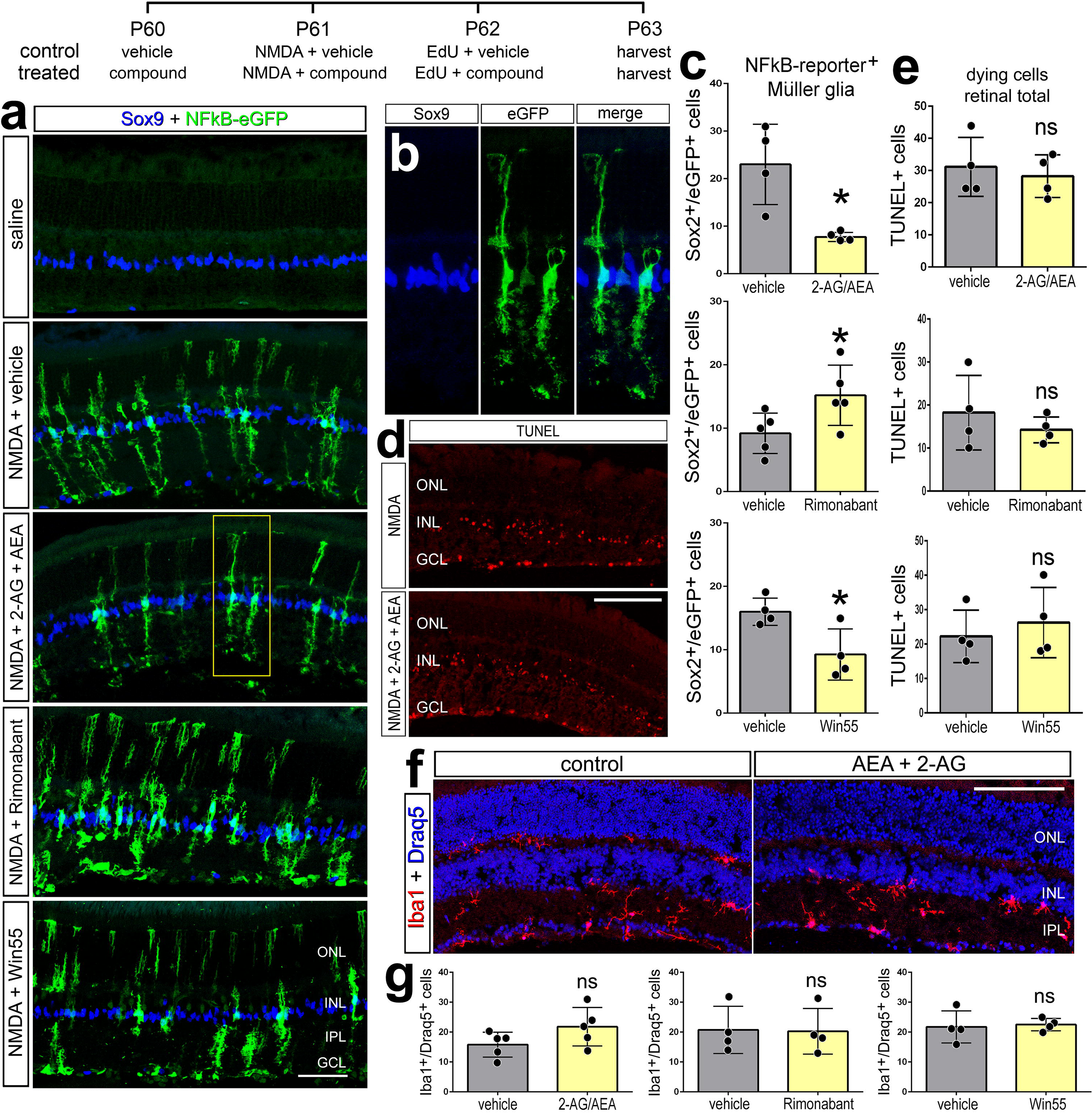
eCBs and NF-kB-signaling in MG of damaged mouse retinas. The treatment paradigm is illustrated at the top of the figure. Eyes of mice (cis-NF-kB^eGFP^) were pretreated with compounds or vehicle prior to NMDA + vehicle/compound, and retinas harvested 24 hrs after the last injection. Compounds included 2-AG+AEA (CNR1 agonists), Win55 (CNR1 agonist), and Rimonabant (CNR1 antagonist). Retinal sections were labeled for Sox9 (blue) and eGFP (green) (**b**), fragmented DNA using the TUNEL method (**d**), and Iba1 (red) and Draq5 (blue; **f**). The histogram/scatter-plots illustrate the mean (±SD) number of eGFP+ MG (**c**), dying cells (**e**) or Iba1+/Draq5+ cells (**g**). Each dot represents one biological replicate. Significance of difference (*p<0.05) was determined by using a paired *t*-test. The calibration bars panels **a**, **c, e,** and **g** represent 50 µm. Abbreviations: ONL – outer nuclear layer, INL – inner nuclear layer, IPL – inner plexiform layer, GCL – ganglion cell layer.

**Figure 9.**
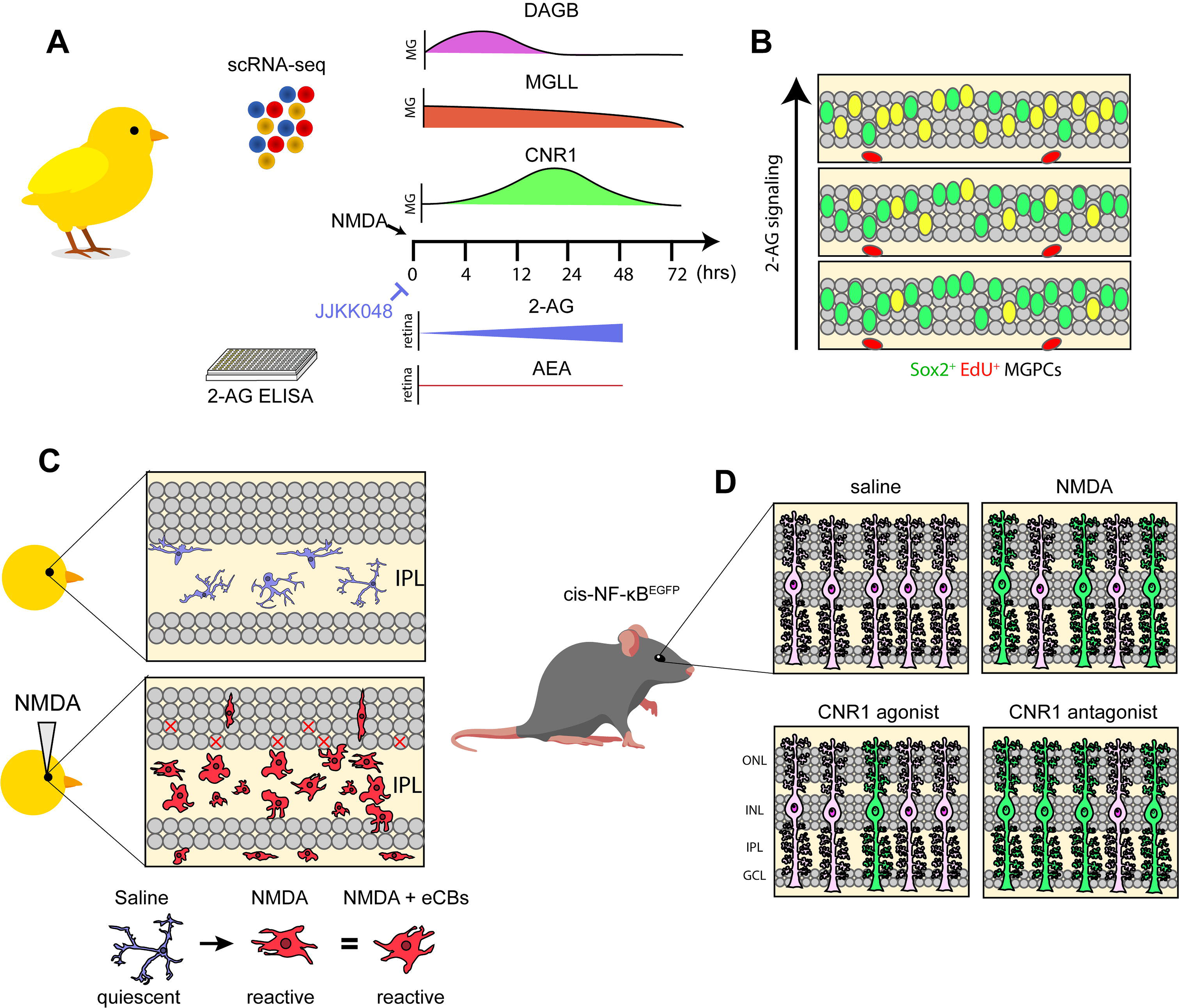
Summary Schematic of eCB effect on MG reprogramming. Using scRNA-seq analysis, we identified patterns of expression of genes involved in eCB synthesis, degradation and signaling (**a**). ELISAs indicated a more prominent role of 2-AG over AEA, and drugs which elevated 2-AG signaling increased MGPCs after damage (**b**). Microglia were unresponsive to these treatments and retained a reactive phenotype in a damaged retina (**c**). In the NF-kBeGFP+ reporter mice, damage activates signaling in MG, which is decreased by exogenous eCBs or CNR1 agonists (**d**).

## Discussion

In this study we investigated the roles of eCB-signaling in the chick model of MG reprogramming. Retinal cells widely expressed both *CNR1* and genes involved in the synthesis and degradation of eCBs. The levels of expression and proportion of MG that express these genes significantly change following damage and during the transition to a proliferating progenitor-like cell. These changes in expression imply functions for eCBs in damaged retinas and during the formation of MGPCs. Indeed, we found that reprogramming of MG into proliferating MGPCs was promoted by eCBs and by CNR1 agonists or enzyme inhibitor that increase retinal levels of 2-AG. Microglia maintained a reactive phenotype in damaged retinas regardless of treatment with eCB drugs. These findings support recent reports that the inflammatory state of MG is important to the transition from resting to reactive, and then to a progenitor-like cell (Fischer et al., 2014; Hoang et al., 2020; White et al., 2017).

### eCB signaling gene expression

In the embryonic chick retina the expression of CNR1 and MGLL has been reported in MG (da Silva Sampaio et al., 2018). In addition to expression in MG, we detected *CNR1* in MG and in a population of amacrine cells and MGLL was detected in MG, some types of ganglion cells and oligodendrocytes. The eCB-related genes were present in MG but at low levels in a small proportion of MG. This pattern of expression is in contrast with high-expressing glial markers, such as glutamine synthetase (*GLUL*), retinaldehyde binding protein 1 (*RLBP1*) and carbonic anhydrase 2 (*CA2*) that are detected in >96% of MG in scRNA-seq preparations (Campbell et al., 2021; Palazzo et al., 2020). We believe this may be due to sensitivity limitations of the reagents wherein low-copy transcripts may not be readily detected. It is also possible that a sub-population of MG express eCB-related genes, suggesting heterogeneity among MG types. However, the eCB-expressing MG subsets are scattered homogenously in these clusters and do not correlate with unique markers corresponding to biologically unique subclusters.

### Elevated eCBs promote MG reprogramming

Although the roles of eCBs have been investigated in the visual system, little is known about how eCBs influence MG reprogramming in different models of retinal regeneration. We observed that exogenous eCB increased the proportion of MG that formed proliferating MGPCs. This effect was reproduced by inhibition of MGLL with JJKK048 which is expected to increase retinal levels of 2-AG. Similarly, this drug has been validated to target MGLL and increase levels of 2-AG levels in mice (Hillard, 2015). Orlistat has been shown to inhibit DAGL and suppress 2-AG synthesis in humans (Bisogno et al., 2006). Although we observed a decrease in number of proliferating MGPCs with Orlistat treatment, we did not observe a significant decrease in 2-AG as measured by ELISA. This may have resulted from the low sensitivity threshold for detecting 2-AG. These lipids represent a very small fraction of total lipids from whole-retina extracts. Alternatively, Orlistat could be targeting fatty acid synthase (FASN), disrupting lipid metabolism to influence retinal levels of 2-AG (Kridel et al., 2004).

We examined whether CNR1 may have mediated eCB effects applying selective agonists and antagonists, drugs with validated specificity in the chick CNS (Ádám et al., 2008; Stincic and Hyson, 2011). We observed complimentary effects with Win55 (CNR1 agonist) increasing and rimonabant (CNR1 antagonist) decreasing numbers of proliferating MGPCs. Nevertheless, we cannot exclude the possibility that these effects are due to indirect actions at MG given that amacrine and ganglion cells express significant levels of *CNR1* and may have mediated effects on MGPCs through secondary factors. We failed to detect *CNR1* expression among microglia, NIRG cells or oligodendrocytes in normal retinas or after NMDA-treatment.

Our findings support the hypothesis that MG are receptive and responsive to eCBs. In different animal models and cell types changes in cell physiology are mediated via interactions and cross-talk with other cell-signaling pathways, such as Notch1 (Frampton et al., 2010), mTor (Palazuelos et al., 2012), MAPK/PI-3K (Dalton et al., 2009), and Wnt signaling (Nalli et al., 2019). These cell-signaling pathways are known to be active and promote the reprogramming of MG into MGPCs in the chick model (Fischer et al., 2002; Gallina et al., 2016; Ghai et al., 2010; Zelinka et al., 2016). However, we have yet to identify the interactions between eCB-signaling and other cell-signaling pathways that have been implicated in the reprogramming of MG. These connections may be difficult to identify in undamaged retinas given that homeostatic enzymes reduce eCB levels and because the sensitivity of MG to eCBs may increase after damage with increased prevalence of *CNR1-*expression among MG.

### eCBs are not neuroprotective to excitotoxic NMDA damage

eCBs have been shown to provide neuroprotection in degenerative retinal diseases (Rapino et al., 2018). Recent articles have even suggested that 2-AG can mediate neuroprotection against AMPA toxicity in the rat retina (Kokona et al., 2021). We investigated eCB-related neuroprotection because levels of retinal damage and cell death are known to influence the reprogramming of MG in to MGPCs. Although, injections of 2-AG and AEA did not impact numbers of dying cells, the CNR1 agonist Win55 increased cell death in the chick retina (supplemental Fig. 2a,b). This could result from interactions with ion channels that are known to occur with these lipophilic eCB ligands (Pertwee, 2010). Alternatively, differences in excitotoxicity with eCB administration could be due to targeting NMDA vs AMPA receptors in the damage model. (2021) reported the death of photoreceptors with AMPA-selective agonists, which does not occur with NMDA. In other disease models where 2-AG provides neuroprotection the mode of cellular damage is slow and progressive (Centonze et al., 2007), unlike our model of NMDA-induced excitotoxicity which acute and severe.

### Microglia reactivity is not influenced by eCBs

eCBs are believed to be potent anti-inflammatory drugs in the CNS (Ullrich et al., 2007). This is frequently suggested as mechanism of clinical benefit in pathological states. In the chick model of reprogramming, we have used dexamethasone GCR receptor agonist to repress the reactivity of microglia (Gallina, 2015). Similarly, treating damaged retinas with NF-kB inhibitor sulfasalazine also resulted in a decrease in the reactive proliferation of CD45^+^ cells (Palazzo et al., 2020). With eCBs and small molecule drugs, there was no evidence that the reactivity of the microglia was influenced. Studies have demonstrated that reduced accumulation of reactive microglia results in neuroprotective effects whereas increased accumulation of microglial reactivity can be detrimental to neuronal survival (Fischer et al., 2015; Todd et al., 2019) (Fischer et al., 2105 Glia; Todd et al., 2019 J Neuroinflam; other refs). In the current study, however, inhibition of NAPEPLD increased the accumulation of reactive microglia, while numbers of dying cells were reduced. This may have resulted from multiple cellular targets being directly affected by the NAPEPLD inhibitor since *NAPEPLD* was detected in microglia, MG and inner retinal neurons.

Our findings are consistent with the notion that eCB-signaling is, in part, manifested through MG. However, we cannot exclude the possibility of eCBs mediate changes in production pro-inflammatory cytokines from reactive microglia in damaged retinas. The relationship between these inflammatory factors and MGPC formation is complex and time-dependent. For example, decreased retinal inflammation from inhibition of microglial reactivity with glucocorticoid agonists reduced MGPC formation, whereas decreased retinal inflammation from inhibition of NFkB-signaling increased MGPC formation (Gallina, 2015; Palazzo et al., 2020). However, the impact of NFkB-signaling on the formation of MGPCs was reversed when the microglia were ablated (Palazzo et al., 2020). Pro-inflammatory factors likely directly influence MG, with evidence that MG activate NF-kB-signaling and express cytokine receptors in damaged retinas (Palazzo et al., 2020). Further studies are required to determine the impact of pro-inflammatory signals on microglia and MG in eCB-treated retinas to better characterize the coordination between these glial cells.

### eCBs repress NF-kB in mouse MG

While reporter lines for MG do not exist in the chick model, the mouse model of retinal damage was applied to the cis-NF-kB^eGFP^ reporter line to identify cells where p65 translocates into the nucleus and drives the expression of the eGFP-reporter. We find that MG are the primary cell type that activates NF-kB-signaling in NMDA-damaged retinas. This supports prior findings in chick that MG respond to proinflammatory cytokines such as TNF associated with NF-kB signaling (Palazzo et al., 2020). After damage eCBs reduced numbers of GFP^+^ MG, suggesting that eCBs limit the activation of NF-kB in support cells. NF-kB has been implicated as an important pathway in mouse retina that may mediate a switch between reactive gliosis and resting MG (Hoang et al., 2020). Recent studies focused on MG reprogramming have highlighted the importance of the interactions between microglia and MG. The ablation of microglia with CSF1R inhibitor have a dramatic impact on the neurogenic capacity of MG that overexpress Ascl1 (Todd et al., 2020). The absence of microglia induced transcriptomic changes in MG which included the repression of gliosis-associated genes (Todd et al., 2020). The context, timing and specific cell-signaling pathways that influence the reprogramming capacity of MG in the mammalian retina requires further investigation.

## Conclusions

In this study we investigated the impact of eCBs on retinal inflammation and MG reprogramming in the chick model. We found transcriptomic evidence of eCB genes expressed by MG and the expression of these genes was dynamic following injury and during the transition into MGPCs. Increasing levels of eCBs through intravitreal injections or upregulation of 2-AG via enzyme inhibitors increased numbers of proliferating MGPCs. Surprisingly, cell death and microglia reactivity were largely unaffected by experimental manipulation of levels eCBs in both chick and mouse models of retinal damage. These data support recent evidence that inflammatory signaling play a pivotal role in regulating reactive gliosis, promoting the de-differentiation in MG, and suppressing the neurogenic capacity of MGPCs.

## Author contributions

WAC – experimental design, execution of experiments, collection of data, data analysis, construction of figures and writing the manuscript. SB and AR - execution of experiments, collection of data and data analysis. TH and SB facilitated the single cell experiments. AJF – experimental design, data analysis, construction of figures and writing the manuscript.

## Data availability

RNA-Seq data are deposited in GitHub https://github.com/jiewwwang/Single-cell-retinal-regeneration

https://github.com/fischerlab3140/scRNAseq_libraries

scRNA-Seq data can be queried at https://proteinpaint.stjude.org/F/2019.retina.scRNA.html.

## Acknowledgements

This work was supported by RO1 EY022030-08, RO1 EY032141- 01 (AJF) and UO1 EY027267-04 (AJF).

