## Supplemental Tables and Figures for "Cannabinoid signaling promotes the reprogramming of Muller glia into proliferating progenitor cells"

**Supplemental Figures**

**Supplemental Figure 1: UMAP-clustering of MG from control and NMDA-treated retinas.** scRNA-seq was used to identify patterns of expression of MG- and MGPC-related genes. Data are illustrated presented in UMAP plots (**a**-**h**) and violin plots (**i**). scRNA-seq libraries from control and treated 3hr, 12hr, and 48hr after NMDA-treatment (**b**). UMAP plots include aggregates of all retinal cells (**a**-**c**) or MG that were re-embedded for UMAP-ordering (**d**-**h**). UMAP-ordered cells formed distinct clusters of resting MG, early activated MG, activated MG and late activated MG + MGPCs (**c-e**). UMAP heatmaps of *CNR1*, *MGLL, NAPEPLD, DAGLA, DAGLB* and *FAAH* demonstrate patterns and levels of expression across different retinal cells, with black dots representing cells with expression of 2 or more genes (**f**-**h**). Violin plots illustrate relative levels of expression in MG from retinas treated with saline, or 3hrs, 12hrs and 48hrs after NMDA-treatment (**i**). Violin plots illustrate percent expressing cells and levels of gene expression and significant changes (*p<0.01, **p<10exp-10, ***p<10exp-20) in levels that were determined by using a Wilcox rank sum with Bonferroni correction.

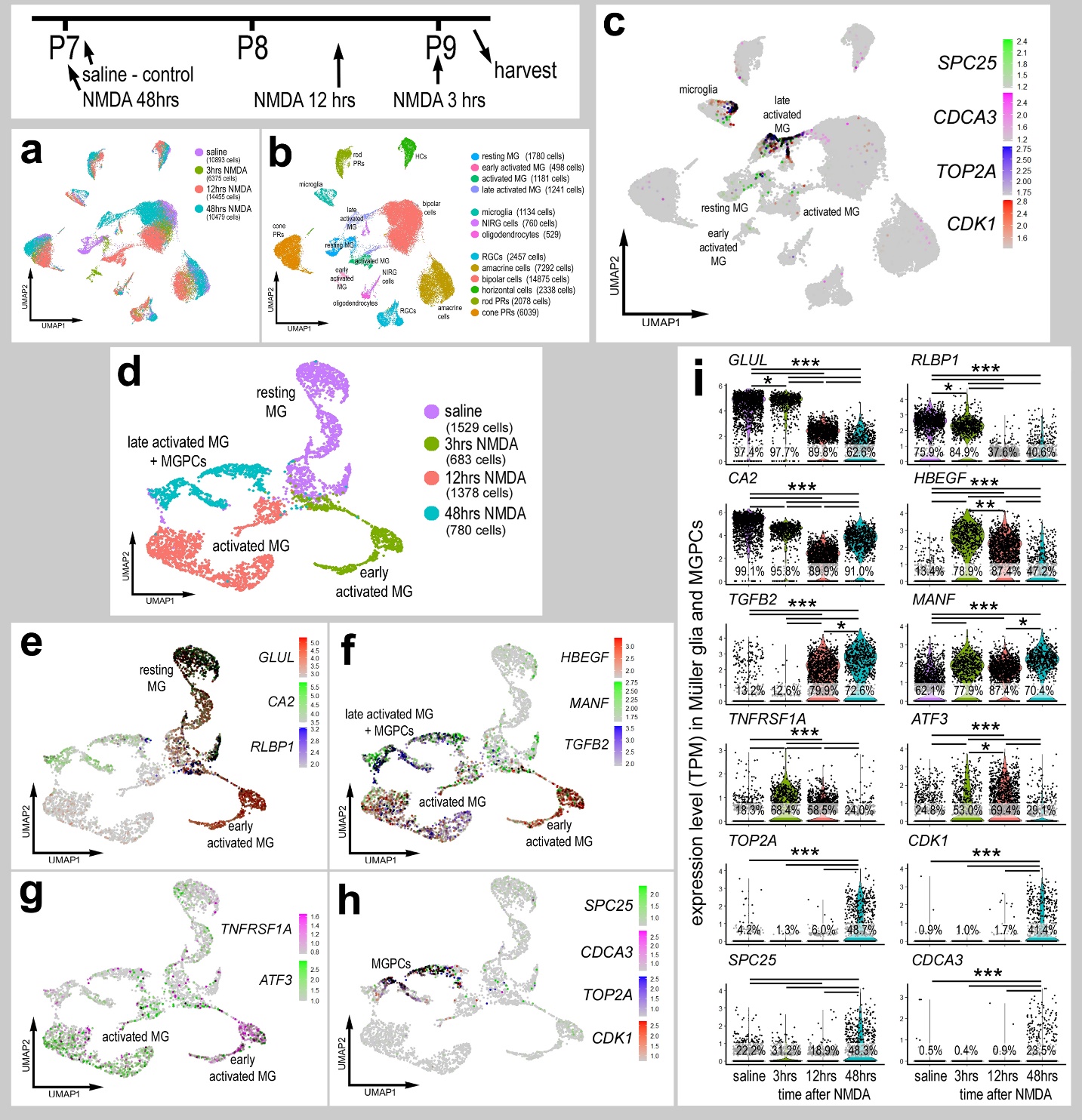

**Supplemental Figure 2. CNR1 agonist increases cell death in NMDA-damaged chick retinas.** The treatment paradigm is illustrated at the top of the figure. Numbers of TUNEL^+^ positive cells (red; **a**) were significantly increased when CNR1 agonist (Win55) was applied. The histogram/scatter-plot in **b** illustrates the mean (±SD) number of labeled cells. Each dot represents one biological replicate. Significance of difference (*p<0.05) was determined by using a paired *t*-test. The calibration bars panels **a**, **c, e,** and **g** represent 50 µm. Abbreviations: ONL – outer nuclear layer, INL – inner nuclear layer, IPL – inner plexiform layer, GCL – ganglion cell layer.

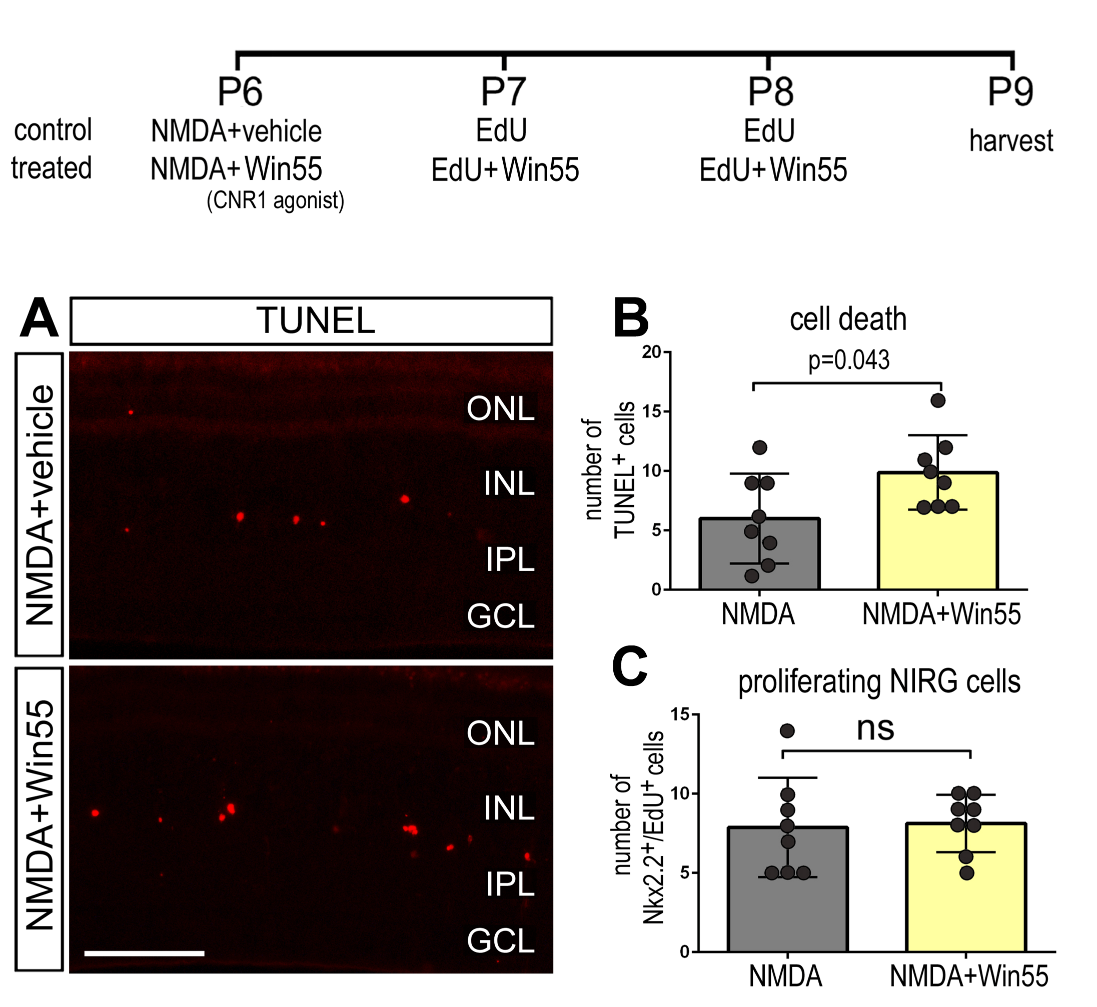
